# IL-21 has a critical role in establishing germinal centers by amplifying early B cell proliferation

**DOI:** 10.1101/2022.01.21.476732

**Authors:** Alexandra R. Dvorscek, Craig I. McKenzie, Marcus J. Robinson, Zhoujie Ding, Catherine Pitt, Kristy O’Donnell, Dimitra Zotos, Robert Brink, David M. Tarlinton, Isaak Quast

## Abstract

The proliferation and differentiation of antigen-specific B cells, including the generation of germinal centers (GC), are prerequisites for long-lasting, high-affinity antibody-mediated immune protection. Affinity for antigen determines B cell recruitment, proliferation, differentiation and competitiveness in the response, largely through determining access to T cell help. However, how T cell derived signals contribute to these outcomes is incompletely understood. Here we report how the signature cytokine of follicular helper T cells, IL-21, acts as a key regulator of the initial B cell response. By activating AKT and S6, IL-21 accelerates cell cycle progression and the rate of cycle entry of B cells, increasing their contribution to the ensuing GC. This effect occurs over a wide range of initial B cell receptor affinities and the resultant increased proliferation can explain the IL-21-mediated promotion of plasma cell differentiation. Collectively, our data establish that IL-21 acts from the outset of a T cell dependent immune response to increase cell cycle progression and fuel cyclic re-entry of B cells thereby regulating the initial GC size and early plasma cell output.

**Summary:** The cytokine IL-21 is a regulator of B cell responses, increasing antibody quantity and quality. Here, we report that during germinal center initiation, IL-21 acts to increase the response magnitude by accelerating cell cycle speed and rate of entry.

## Introduction

The functionality of T cell dependent (TD) B cell responses, which underlie almost all vaccine success, relies on germinal centers (GC). GC are specialized, transient structures located within follicles of secondary lymphoid organs. Here, B cells mutate the genes encoding their antigen receptor (B cell receptor, BCR) with those gaining higher affinity for antigen being selected by their interaction with T follicular helper cells (Tfh) to differentiate into antibody secreting plasma cells, long-lived memory B cells or to undergo further rounds of proliferation and BCR diversification [1, 2]. Within GC, Tfh-derived signals are considered to control B cell proliferation and selection [3-6], while during the initial phase of the response, B cell intrinsic determinants such as BCR affinity and avidity govern response participation [7, 8]. BCR ligation triggers a signaling cascade that influences B cell fate in an antigen affinity-dependent manner [9, 10] including survival, proliferation and differentiation (reviewed in [11]). In addition, naïve B cells capture antigen from the surface of antigen presenting cells using pulling forces, with the BCR affinity determining the efficiency of this process and thus the access to T cell help [12, 13]. The outcome of cognate T:B interaction is then dependent on the expression of co-stimulatory molecules, adhesion molecules and the duration of the T:B interactions [4, 6]. T cell derived cytokines such as IL-4, IL-10, IL-13 and IL-21 can also modulate human and mouse B cell proliferation, apoptosis and differentiation and thus potentially influence GC initiation [14-20]. One of these cytokines, IL-21, is produced by follicular helper T cells shortly after the initiation of a TD B cell response then gradually increases in amount until GC reach maturity [21-24]. The outcome of IL-21 signaling in B cells *in vitro* are multiple and context-dependent, including co-stimulation, growth arrest or apoptosis [25] as well as promoting plasma cell differentiation and supporting antibody class switching [17, 26]. While IL-21 has a key role in maintaining GC [27-29], whether any of its multiple activities contribute to naïve B cell activation and recruitment into the TD B cell response *in vivo* is unresolved. This prompted us to investigate the role of IL-21 during TD B cell response initiation.

## Results

To study the involvement of IL-21 in initiating TD B cell responses, we developed an adoptive transfer and immunization system with defined, identifiable cognate T and B cell partners. WT or *Il21r*^-/-^ mice were crossed with mice that carried a knock-in rearranged BCR specific for hen egg lysozyme (BCR-HEL), known as SW_HEL_ mice [30, 31]. Additionally, these mice were crossed with *eGFP* transgenic mice and all WT or *Il21r*^*-/-*^ SW_HEL_ mice used in this study were *Rag1* deficient, preventing endogenous BCR rearrangement during development and thus unintended B cell activation. In the resultant B cell donor mice, essentially all B cells were specific for HEL (Fig. EV1A) and expressed eGFP (Fig. EV1B). Donor CD4 T cells were derived from mice transgenic for the alpha and beta chains of a CD4 restricted T cell receptor (TCR) specific for ovalbumin, known as OTII [32], and carried a *GFP* knock-in at the *Il21* locus (*Il21*^Gfp/+^), allowing for analysis of *Il21* transcription via GFP fluorescence [22]. As an antigen, we generated a recombinant protein of HEL fused with the I-A^b^-restricted 12-mer peptide recognized by OTII T cells [33], referred to as HEL^WT^OVA_pep_. To study the role of affinity, we introduced 2 or 3 mutations in the sequence encoding HEL, referred to as HEL^2X^OVA_pep_ and HEL^3X^OVA_pep_, resulting in a SW_HEL_ BCR affinity series of 2 × 10^10^ M^−1^ (HEL^WT^), 8 × 10^7^ M^−1^ (HEL^2x^), and ∼1 × 10^7^ M^−1^ (HEL^3X^) [34, 35] (Fig. EV1C).

### IL-21 promotes B cell expansion from the outset of a TD immune response by increasing cell cycle speed and rate of entry

Having established an experimental system, we investigated the role of IL-21 in the response to a moderate affinity antigen, HEL^2X^OVA_pep_. CD45.2 WT and *Il21r*^-/-^, cell-trace violet (CTV) labelled SW_HEL_ B cells (5 × 10^4^ of each) were co-transferred with 5 × 10^4^ *Il21*^Gfp/+^ OTII T cells into CD45.1 congenic recipients and immunized *ip* with alum-adsorbed HEL^2X^OVA_pep_ (Fig. 1A). This setup allowed identification and analysis of WT and *Il21r*^-/-^ SW_HEL_ B cells within the same recipient mouse (Fig. EV1D). All SW_HEL_ B cells had started to proliferate at day 3.5 post immunization, but by day 4.5 the expansion of *Il21r*^-/-^ B cells was reduced significantly compared to their WT counterparts in the same mouse (Fig. 1B).

**Fig. 1:**
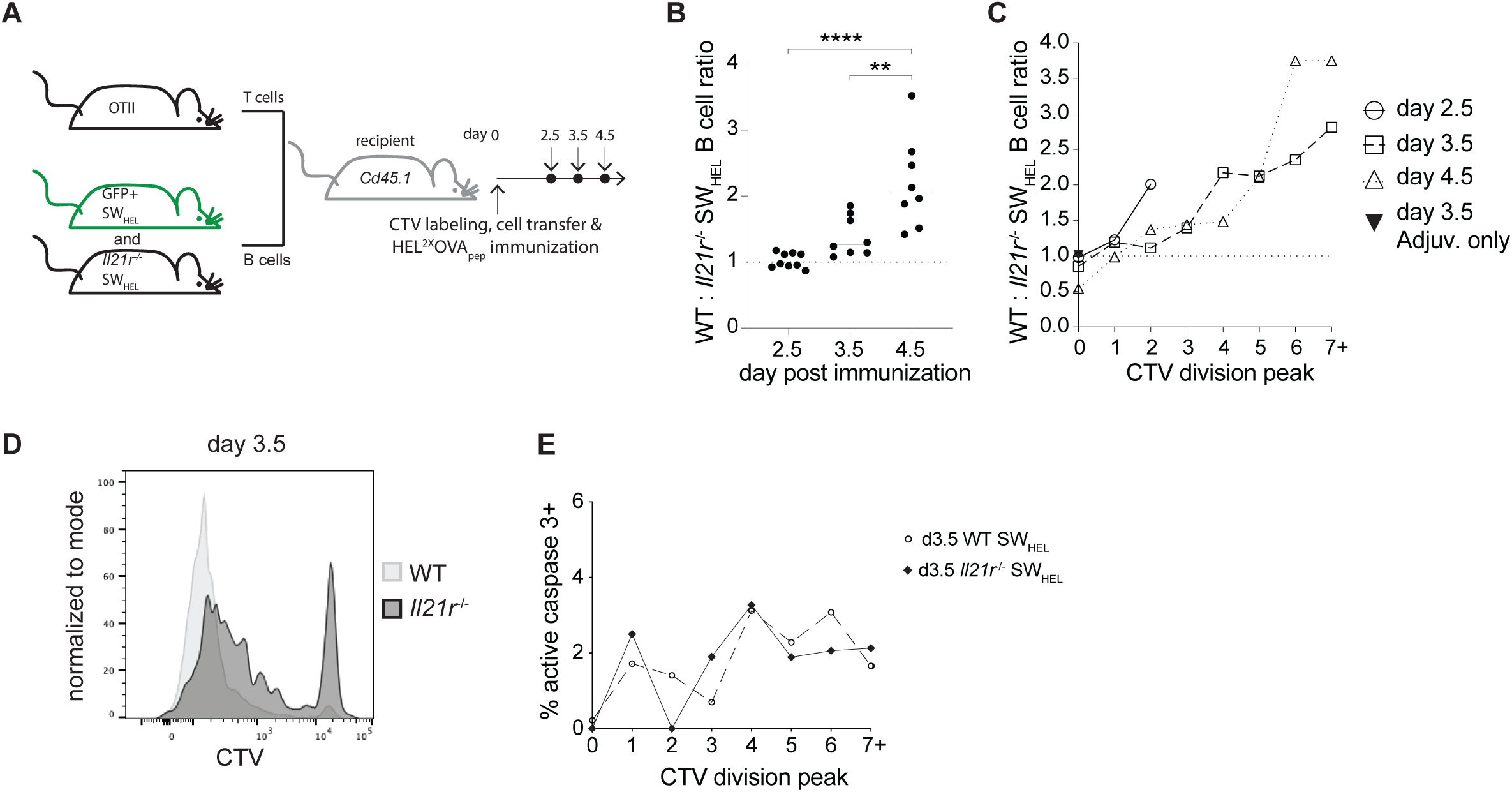
IL-21 promotes B cell expansion. **(A)** Experimental setup to study expansion of WT and *Il21r*^-/-^ SW_HEL_ B cells. **(B)** Ratio of WT to *Il21r*^-/-^ splenic SW_HEL_ B cells over time. **(C)** Cell ratio within CTV division peaks. **(D)** CTV profile on day 3.5. **(E)** Rate of apoptosis measured by detecting active caspase 3 by flow cytometr. C-E show concatenated data from 4-5 mice representative for two independent experiments. Data in B were pooled from two independent experiments with statistical analysis by one-way ANOVA with Tukey’s post-test. ** p ≤ 0.01; **** p ≤ 0.0001.

Cell division analysis by CTV dye dilution revealed that over time, *Il21r*^-/-^ B cells were progressively and increasingly disadvantaged, being less likely to enter into subsequent divisions as indicated by an increasing proportion of *Il21r*^-/-^ cells not further diluting CTV (Fig. 1C, D). IL-21 has been implicated in regulating B cell homeostasis by increasing apoptosis in the absence of CD40 signaling in *in vitro* experiments [36]. To assess if IL-21 regulated apoptosis during early B cell expansion *in vivo*, we analyzed each CTV peak of WT and *Il21r*^-/-^ SW_HEL_ B cells on day 3.5 post HEL^2x^OVA_pep_ immunization for the presence of active caspase 3, indicative of the onset of apoptosis. Active caspase 3 positive cells, although rare, were at a similar frequency within each division peak of WT and *Il21r*^-/-^ SW_HEL_ B cells (Fig. 1E), in line with previous reports [24].

With cell death an unlikely cause for the reduced representation of *Il21r*^-/-^ SW_HEL_ B cells, we investigated whether IL-21 influenced cell cycle progression. To assess this, S phase cells were time-stamped by incorporation of the DNA nucleoside analogue 5-bromo-2’-deoxyuridine (BrdU), which has a short bioavailability with most labelling occurring within 30 min to 1h of injection [37]. To track the subsequent progression through the cell cycle, we analyzed the cells’ DNA content at various time points thereafter, an approach that has been used to dissect cell cycle progression of T cells [38]. Accordingly, cells in active DNA synthesis on day 3.5 post HEL^2x^OVA_pep_ immunization were labeled with BrdU (Fig. 2A) and analyzed 4, 10 and 12h thereafter. In most instances, a higher proportion of WT SW_HEL_ B cells incorporated BrdU compared to *Il21r*^-/-^ SW_HEL_ B cells in the same animal, confirming that IL-21 increased cell cycle activity (Fig. 2B). To determine if IL-21 enhanced the rate of cell cycle entry, the speed of cell cycle transition or a combination of both, DNA content of BrdU^+^ cells 4, 10 and 12h post BrdU injection was measured by co-staining *ex vivo* with 7-Aminoactinomycin D (7-AAD). Rapidly proliferating B cells have been measured to complete a cell cycle within 8-12h of initiation [39] with S phase comprising 5-6h [5], correlating closely with total cell cycle time [39]. Therefore, assessing the proportion of BrdU^+^ cells that had completed cell division (that is had 2N DNA content) 4h after BrdU injection allowed us to determine the relative speed of S phase completion and to do so independently of the rate of cell cycle re-entry, as only cells that had completed one S phase were analyzed (Fig. 2C, left). This revealed that the proportion of BrdU^+^ cells with 2N DNA content, indicative of the cells having progressed from S to G1 within 4 hours, was significantly higher in WT than *Il21r*^-/-^ SW_HEL_ B cells, indicating more rapid cell cycle progression (Fig. 2D). Ten hours post BrdU injection, twice the time required to complete S phase, a 2N DNA content in BrdU^+^ cells identified cells that had divided once but not re-entered S phase in a subsequent cell cycle (Fig. 2C, right). The reduced proportion of BrdU^+^ cells with a 2N DNA content for WT SW_HEL_ B cells compared to *Il21r*^-/-^ B cells indicated a greater fraction of WT SW_HEL_ B cells had re-entered division after 10h, resulting in more cells in S/G2 (Fig. 2D). After 12h, enough time for some cells to have completed 2 rounds of division, 2N DNA content of BrdU^+^ cells in both genotypes was proportionately similar (Fig. 2D).

**Fig. 2:**
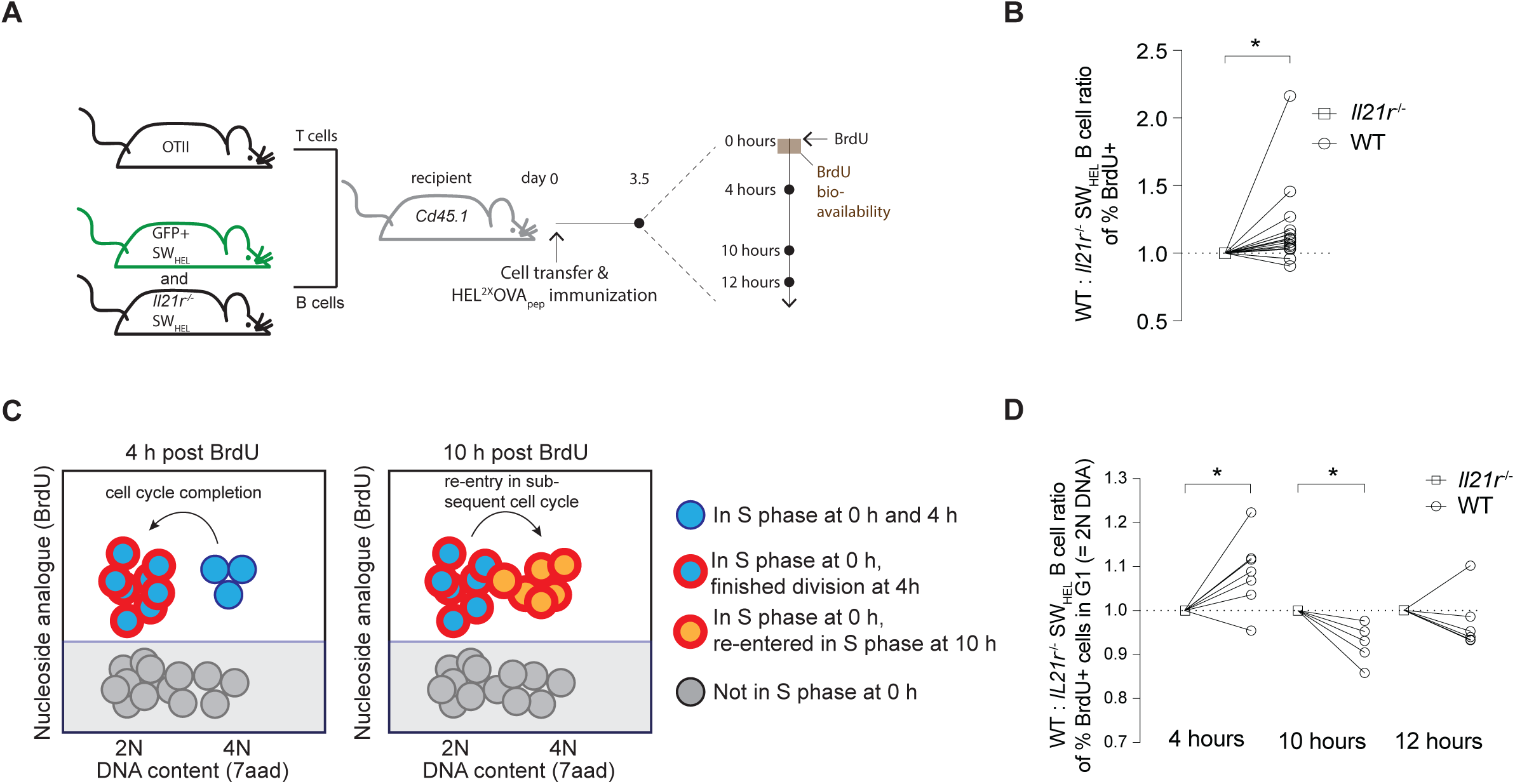
IL-21 increases cell cycle speed and rate of re-entry. **(A)** Experimental setup of BrdU pulse on day 3.5 post immunization. **(B)** WT to *Il21r*^-/-^ SW_HEL_ B cell ratio of cells that have been in S phase and thus incorporated BrdU (pooled data from 4, 10 and 12h time points). **(C)** Schematic depiction of BrdU and DNA content (7AAD) analysis. At 4h, BrdU positive cells with 2N DNA content mark those that have finished the cell cycle, whereas at 10h 2N DNA content identifies cells that have not yet entered the subsequent cell cycle. **(D)** Ratio of the proportion of BrdU positive WT and *Il21r*^-/-^ SW_HEL_ B cells with 2N DNA content on day 3.5 post immunization at 4, 10 and 12h post BrdU pulse. Data were pooled from two independent experiments. Statistical analysis by one-sample t test. *p ≤ 0.05.

The reduced re-entry of cycling *Il21r*^-/-^ SW_HEL_ B cells seen 10 h after the BrdU pulse could be due to increased S phase duration and concomitantly increased cell cycle duration, giving the cells less time from completion to re-entry [39]. Equally or additionally, the rate of cell cycle re-entry could be reduced, with the time spent in G1 prolonged, in the absence of IL-21. To investigate if IL-21 influenced the rate of cell cycle initiation, we developed an experimental system in which endogenous IL-21 production and signaling were abrogated (*Il21* and *Il21r* double deficient recipient mice and *Il21*^-/-^ OTII T cells) allowing IL-21 to be provided as a pulse that acted only on the transferred B cells (Fig. 3A). On day 3.5 post cell transfer and immunization, 2 µg of recombinant IL-21 or saline was injected *iv*. Following this, BrdU was injected one hour afterwards to label and exclude cells that had already initiated a cell cycle at the time of IL-21 injection. Consequently, cells that were subsequently BrdU^-^ were presumed to be in G1 at the time of IL-21 injection. Ten hours after IL-21 injection, the DNA content of BrdU^-^ cells was analyzed, with cells having >2N DNA content being those that had entered S phase approximately 2-5 hours following IL-21 or saline injection (Fig. 3B). This time point was chosen to allow time for B cells to respond to IL-21 stimulation, complete G1 and enter S phase. To determine the rate of cell cycle entry independent of the extent of cell proliferation by WT and *Il21r*^-/-^ SW_HEL_ B cells over time, the proportion of BrdU^-^ cells that had entered S phase (DNA content by 7-AAD >2N) was determined for both genotypes (Fig. EV1E) and the ratio within each mouse calculated. In the absence of IL-21 injection, the ratio of WT to *Il21r*^-/-^ SW_HEL_ B cells that had entered cell division was randomly distributed. In contrast, following IL-21 administration BrdU^-^ cells with >2N DNA content, and thus in S/G2, were more frequent among WT cells in all but one mouse (Fig. 3C). This suggested that promotion of cell cycle entry, in addition to increased S phase speed, was a direct consequence of IL-21 signaling in B cells.

**Fig. 3:**
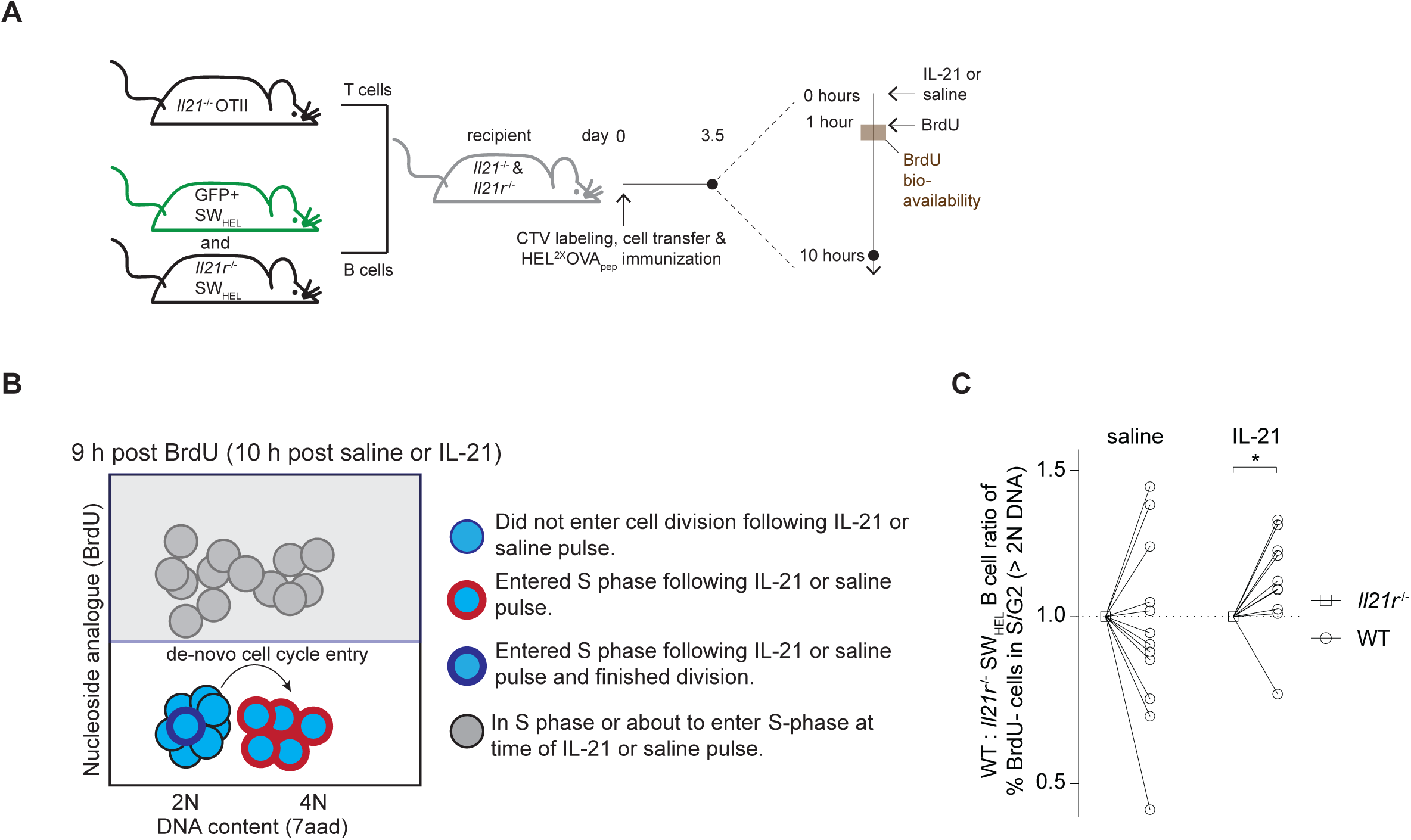
IL-21 directly promotes cell cycle entry. **(A)** Experimental setup of *in vivo* IL-21 pulse. **(B)** Schematic depiction of BrdU and 7AAD analysis. Cells that had not already been in or were about to enter S phase at the time of IL-21 treatment were identified as being BrdU negative. > 2N DNA content among BrdU negative cells marked those that have entered the cell cycle following saline or IL-21 treatment. **(C)** Rate of *de novo* cell cycle entry 10h post *in vivo* IL-21 (2 µg) or saline pulse. The ratio of the fraction of WT to *Il21r*^-/-^ cells not in S/G2 at the time of pulse (BrdU^-^ cells) and containing >2N DNA content 10h after pulse is shown. Data were pooled from two independent experiments. Statistical analysis by one-sample t test. * p ≤ 0.05.

### IL-21 synergizes with BCR and CD40 to promote AKT and S6 phosphorylation

IL-21 alone does not initiate B cell proliferation, but B cell proliferation occurs in the absence of IL-21, indicating that IL-21 amplifies rather than initiates mitogenic signals such as those downstream of BCR and CD40 stimulation. Phosphorylation and activation of AKT, a key event downstream of both BCR and CD40 ligation, can lead to the phosphorylation of mammalian target of rapamycin complex 1 (mTORC1) with the subsequent activation of S6-kinase (S6K) [40], which in turn regulates key mediators of cell proliferation including phosphorylation of S6 (p-S6). Importantly, IL-21R signaling has been shown to result in AKT phosphorylation (p-AKT) at serine 473 (S473) in cell lines and CD8 T cells [41]. To explore the possibility that IL-21 promoted the cell cycle via enhancing AKT and S6-phosphoryation, we incubated splenocytes *ex vivo* with or without 20 ng IL-21 for 3 h in the presence of agonistic anti-CD40, also for 3h, or BCR stimulation using biotinylated anti-IgK/λ followed by streptavidin-mediated cross-linking for the last minute of incubation. Phosphoflow analysis of naïve B cells (Fig. EV2) showed incubation with IL-21 increased AKT phosphorylation at S473, which was further increased by concurrent stimulation through BCR or CD40 (Fig. 4A, B). IL-21R dependency was confirmed with B cells from *Il21r*^-/-^ mice, which retained AKT phosphorylation in response to BCR or CD40 stimulation but were unaffected by exposure to IL-21 (Fig. 4A, B). Phosphorylation of S6 was minimally induced following BCR stimulation and more so following CD40 ligation but potently increased by the presence of IL-21 (Fig. 4C). p-S6 amounts were distributed bimodally with the frequency of p-S6 positive cells increased by exposure to IL-21, and further again by addition of BCR or CD40 signals (Fig. 4D), effects that also required IL-21R expression (Fig. 4C, D). In addition to increasing the proportion of p-S6 positive cells, IL-21 increased the median amount of p-S6 among p-S6 positive cells (Fig. 4E).

**Fig. 4:**
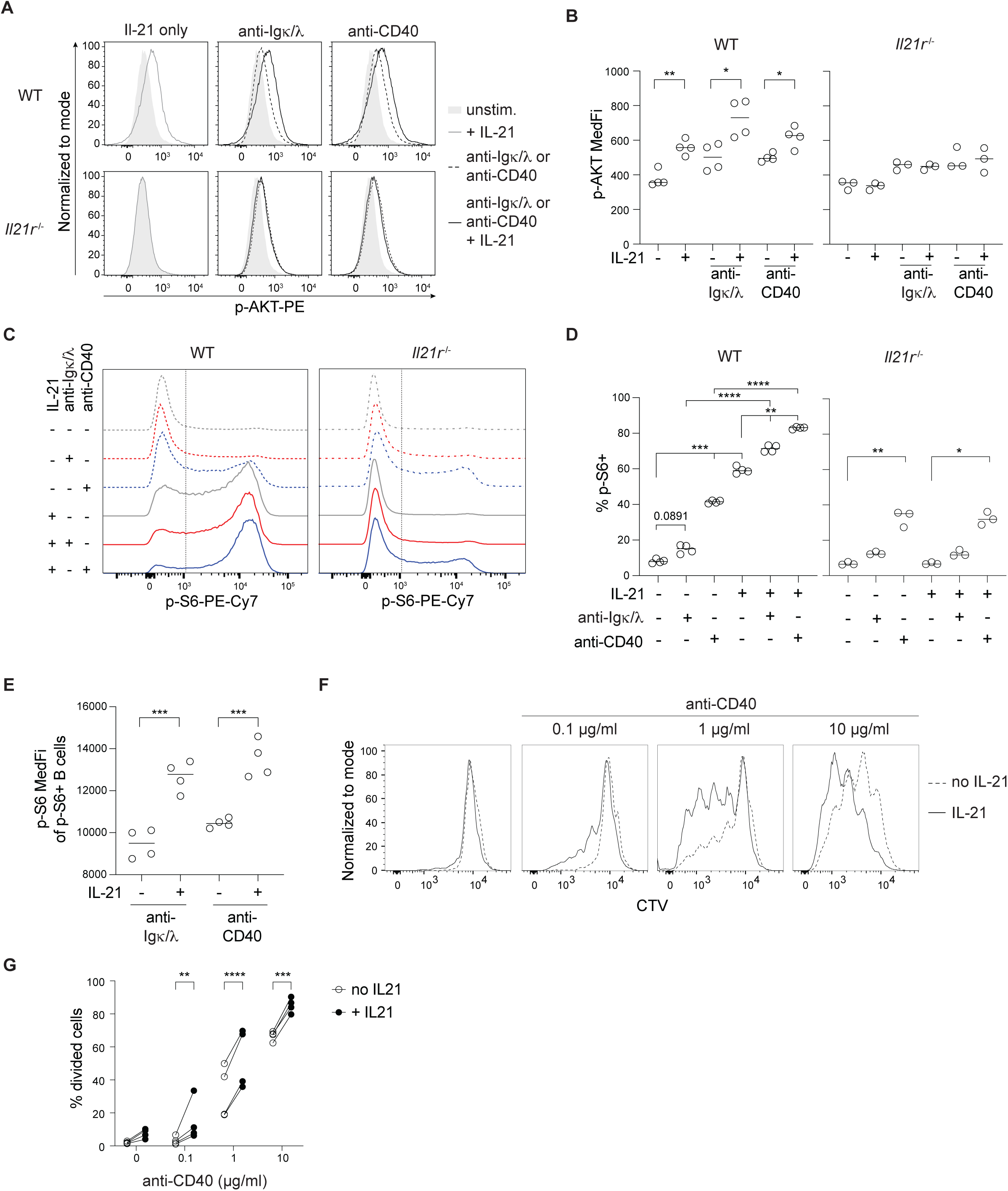
IL-21 synergizes with BCR and CD40 to promote AKT and S6 phosphorylation. **(A-E)** Phosphoflow analysis of naïve WT or *Il21r*^-/-^ B cells following *in vitro* culture for 3h with or without IL-21 (20 ng/mL) and/or in the presence of BCR cross-linking (biotinylated anti-IgK + anti-IgA and avidin-mediated cross-linking) or agonistic anti-CD40. **(A)** Exemplary p-AKT (S473) staining and **(B)** quantification of median fluorescence intensity (MFI). **(C)** Exemplary p-S6 (Ser235/236) staining and **(D)** quantification of frequency of p-S6 positive cells and **(E)** p-S6 MFI of p-S6 positive cells. **(F)** Exemplary CTV division peaks and **(G)** statistical analysis of SW_HEL_ B cells after *in vitro* culture for three days with or without anti-CD40 and IL-21 (20 ng/mL). Individual dots represent biological replicates with data representative of (A-E) or pooled from (G) two independent experiments. Statistical analysis by one-way ANOVA with Tukey’s post-test (D), t test (B, E) or two-way ANOVA with correction for multiple comparison using Šidák method (G). * p ≤ 0.05; ** p ≤ 0.01; *** p ≤ 0.001; **** p ≤ 0.0001.

T cell help via CD40 ligation is essential for TD immune responses *in vivo [42]*, and B cell access to T cell help - and thus CD40 signaling - is determined in large part by BCR affinity [12, 43]. The additive effect of IL-21 on B cell signaling suggested that IL-21 could have influenced this relationship by adjusting the minimal CD40 signaling threshold and/or by amplifying the response to CD40 ligation. To test this, we cultured CTV labeled naïve SW_HEL_ B cells alone or with 10, 1, or 0.1 µg anti-CD40 and in the presence or absence of IL-21 (20 ng/mL) and then analyzed CTV dilution three days later. As expected, the initiation of proliferation was dependent on CD40 signaling in a dose-dependent manner (Fig. 4F, G). Addition of IL-21 increased the extent of CTV dilution such that at the lowest concentration of anti-CD40, CTV dilution occurred only in the presence of IL-21 (Fig. 4F, G). Collectively, these results indicated that IL-21 amplified BCR and CD40 signaling and had lowered the minimal requirement for cell cycle initiation.

### IL-21 amplifies B cell participation in the GC across a wide range of initial BCR affinities

The increased initiation and expedited progression through division in response to IL-21 and the amplification of p-S6 signaling suggested a role for IL-21 in the recruitment of B cells into GC responses. Moreover, with T cell help to B cells being determined by the BCR affinity-dependent efficiency of antigen capture and presentation in the context of MHC-II [12, 43], the increased cell cycle initiation at low anti-CD40 concentrations in the presence of IL-21 *in vitro* suggested that IL-21 may have influenced the entry and continued participation of B cells in a BCR affinity-dependent manner. To investigate the contribution of IL-21 and BCR affinity to B cell recruitment *in vivo*, we immunized mice with HEL^WT^-, HEL^2X^- or HEL^3X^-OVA_pep_ adsorbed on 45 µg alum adjuvant and analyzed the SW_HEL_ B and OTII T cell response 4.5 days post immunization (Fig 5A). For the lowest affinity antigen (HEL^3x^OVA_pep_) we also included a group receiving 90 µg of alum to further boost the response. The number of *Il21*^Gfp/+^ OTII cells that differentiated into CXCR5^+^ PD-1^high^ Tfh cells (Fig. EV3A), was largely independent of the HEL-OVA_pep_ variant used with a statistically non-significant tendency towards higher cell numbers if 90 µg adjuvant was used (Fig. 5B). Similarly, the proportion of *Il21*^Gfp/+^ OTII Tfh cells expressing GFP, and thus transcribing the *Il21* locus, was comparable (Fig. 5C). Thus, early Tfh differentiation and IL-21 production were largely independent of the HEL-OVA_pep_ antigen variant used, effectively creating an experimental system in which the presence or absence of IL-21 signaling to B cells, BCR affinity and adjuvant dose were the only variables. Cell division was monitored by CTV dye dilution of SW_HEL_ B cells, done using spectral cytometry that increased resolution to 11 divisions (Fig. EV3B, C and Fig. 5D). SW_HEL_ B cells proliferated less in the absence than in the presence of IL-21R signaling with HEL^3X^OVA_pep_ immunization such that some CTV dilution was only detected if 90 µg alum was used (Fig. 5D). While we intended to assess the role of IL-21 in B cell response participation by comparing the number of WT and IL-21R deficient SW_HEL_ B cells that remained undivided, few events were obtained and the number varied between mice too much to allow for accurate comparison. To quantify the distribution of rare antigen-specific B cells early during the response, we therefore pooled CTV peaks 0-4, 5-8 and 9-11+ as undivided/minimally proliferative, moderately proliferative and highly proliferative cells, respectively. The number of cells in early peaks was largely unchanged between WT and IL-21R deficient B cells, but a significantly higher proportion of WT SW_HEL_ B cells were present in peaks 9-11+ following HEL^WT^OVA_pep_ and HEL^2x^OVA_pep_ immunization (Fig. 5E). In contrast, HEL^3x^OVA_pep_ immunization resulted in very few mice containing B cells that had undergone more than 8 divisions. In agreement with results in Figure 1, these data indicated that the magnitude of the proliferation deficit caused by IL-21R deficiency increased the more the cells divided. To assess if IL-21 influenced the phenotype of responding B cells, we analyzed CD38, IgD and Fas expression, all known to be dynamically regulated during early B cell activation [44]. As shown in Figure 5F, WT and IL-21R deficient B cells had a highly similar expression profile that was also closely associated with the extent of CTV dilution. CD38 and IgD expression were lost gradually upon consecutive cell divisions while Fas expression increased (Fig. 5F). Early plasma cell differentiation was infrequent, occurred only upon HEL^WT^OVA_pep_ immunization, and was again comparable between WT and IL-21R deficient cells (Fig. 5G). Thus, IL-21 promoted early B cell proliferation without affecting their differentiation.

**Fig. 5:**
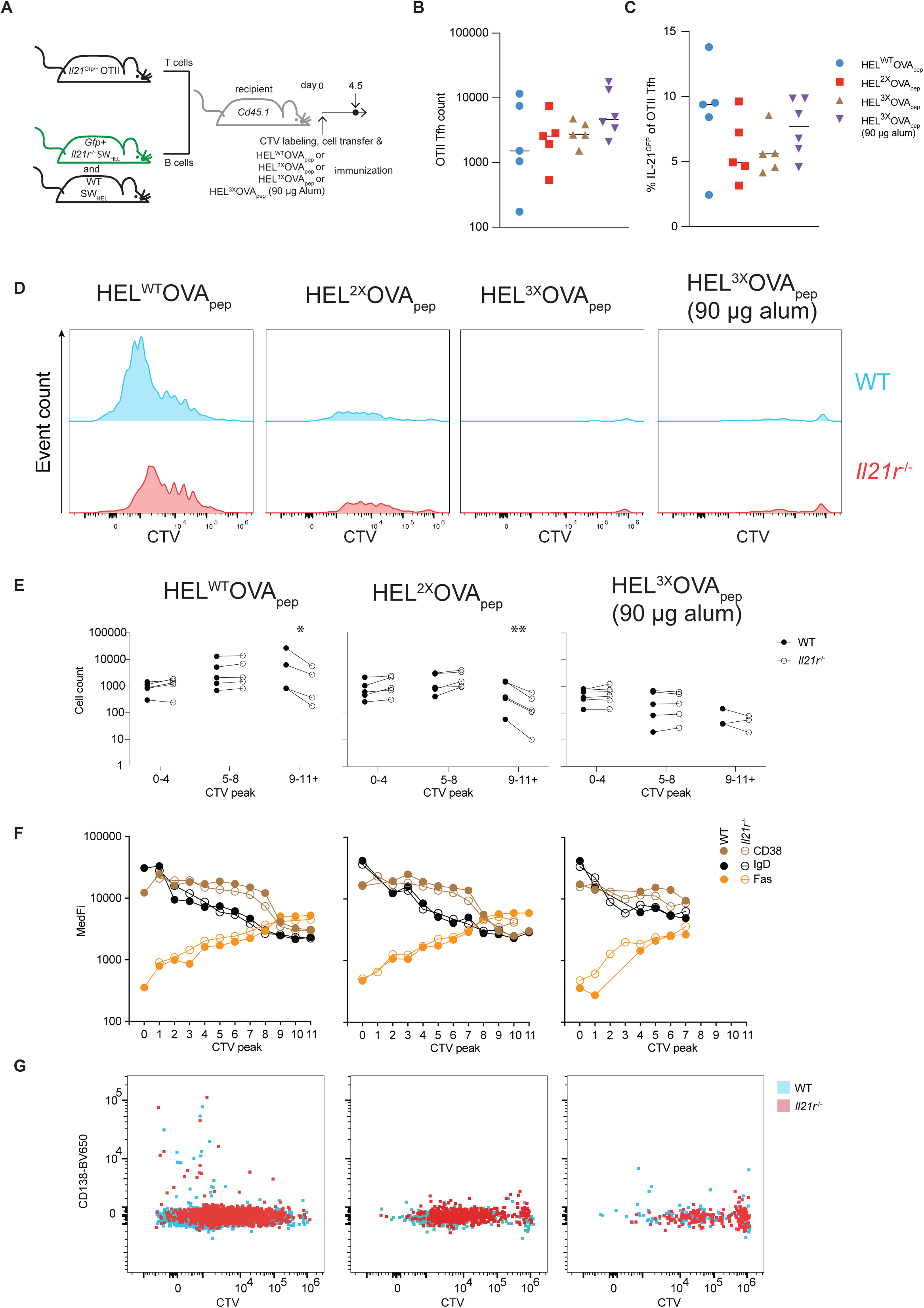
IL-21 amplifies B cell recruitment into the GC. **(A)** Setup to study affinity-dependent effects of IL-21. **(B)** Splenic OTII Tfh count and **(C)** *Il21* locus transcription. **(D)** WT and *Il21r*^-/-^ B cell analysis showing CTV profile and **(E)** absolute cell count in pooled division peaks. **(F)** Cell surface expression of CD38, IgD and FAS in individual division peaks. **(G)** Plasma cell differentiation identified by CD138 expression. D, F and G show concatenated data from 3-5 mice per group. B, C and E show data from two independent experiments with a total of 5-6 mice per group. Statistical analysis by one-way ANOVA with Tukey’s post-test (B, C) or Multiple paired t tests with p values corrected for multiple comparisons using Holm-Šídák method (E). * p ≤ 0.05; ** p ≤ 0.01.

To assess the extent to which the observed effects translated into differences in established GC, we repeated these experiments, this time analyzing on day 7 post immunization. In contrast to day 4.5, SW_HEL_ B cells were now readily detectable in all immunization conditions and showed the canonical FAS^+^ GL7^+^ GC B cell phenotype (Fig. EV4A and B). In response to HEL^WT^OVA_pep_ immunization, the proportion of *Il21r*^*-/-*^ SW_HEL_ B cells with a GC phenotype was significantly reduced compared to WT, with all other immunizations showing the same trend (Fig. 6A). Compared to WT, the absolute number of IL-21R deficient GC B cells was reduced upon immunization with each HEL-OVA_pep_ antigen variant (Fig. 6B), with the difference most pronounced upon HEL^WT^OVA_pep_ immunization (Fig. 6 C). The expansion deficit of IL-21R deficient GC B cells was also reflected in an over-representation of cells with the CD86^+^ CXCR4^low^ GC light zone (LZ), centrocyte phenotype [45] (Fig. EV4C, D), as previously reported [29, 46]. SW_HEL_ PC differentiation, identified by CD98 and CD138 expression (Fig. EV4A), was only consistently detected in HEL^WT^- and HEL^2x^-OVA_pep_ immunized mice and the representation of WT and IL-21R deficient SW_HEL_ cells among PC mirrored that within GC (Fig. 6 D, E). In fact, SW_HEL_ GC B cell and SW_HEL_ PC numbers were closely correlated independent of genotype or affinity for the immunizing antigen, arguing against IL-21 directly promoting PC differentiation, at least at this stage of the response. Collectively, these data revealed IL-21 as crucial in establishing the initial GC size and demonstrated that promotion of continued proliferation, not differentiation, was the dominant effect of IL-21 at this stage of the response.

**Fig. 6:**
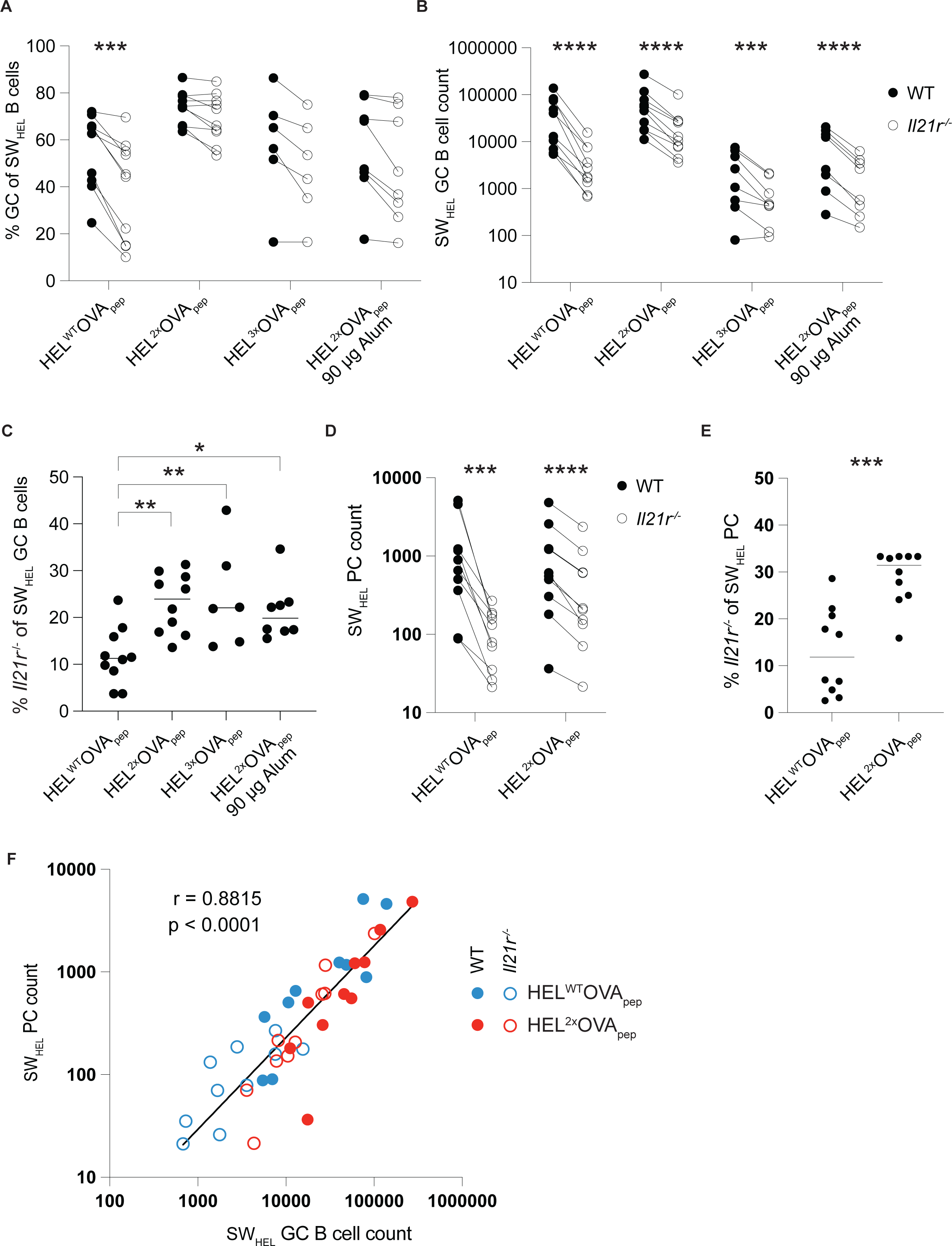
High and low affinity B cell responses are amplified by IL-21. Experimental setup as in Figure 5A but without CTV labeling and analysis on day 7 post immunization. **(A)** Proportion of SW_HEL_ B cells with a FAS^+^GL7^+^ GC phenotype, **(B)** SW_HEL_ GC B cell count and **(C)** representation of IL-21R deficient cells among total SW_HEL_ GC B cells. **(D)** SW_HEL_ PC count and **(E)** representation of IL-21R deficient cells among total SW_HEL_ PC. (**F**) Correlation of SW_HEL_ GC B cell and PC count. Data were pooled from two independent experiments. Statistical analysis by Multiple paired t tests (A, B, D) with p-values corrected for multiple comparisons using Holm-Šídák method or One-way ANOVA with Tukey’s post-test (C) or t test (E). Analysis in F using Pearson correlation coefficient. * p ≤ 0.05; ** p ≤ 0.01; *** p ≤ 0.001; **** p ≤ 0.0001.

## Discussion

The results reported here show IL-21R signaling to be a crucial component of early B cell activation during TD B cell responses, promoting B cell expansion by increasing the speed of cell cycle transition and the rate of entry and re-entry into the cell cycle. These effects are mediated, at least in part, by IL-21 and T cell help additively promoting two key events in triggering cell division, phosphorylation of AKT and S6. Signaling via JAK-STAT, in particular STAT3, is activated downstream of the IL-21R and IL-21R deficiency leads to reduced proliferation and increased apoptosis in the context of CD40 ligation of human B cells [47]. Furthermore, IL-21R signaling results in PI3K and MAPK activation and promotes cell division by inducing S6 phosphorylation (reviewed [48]) and IL-21 was recently reported to promote GC B cell proliferation by sustaining the c-MYC target AP4 [49]. BCR and CD40 employ PI3K and NF-kB pathways to transduce activating signals in B cells that, depending on duration and magnitude, can also lead to the phosphorylation of S6 [40]. While surprisingly little is known about the specific function of S6 in B cells, both AKT and S6 are part of the mTOR signaling network, a key regulator of cell metabolism and proliferation (reviewed [50]). In addition, B cell proliferation, particularly in response to BCR signaling, is highly sensitive to rapamycin, an mTOR inhibitor that also inhibits S6 phosphorylation [51, 52]. Thus, we consider it likely that the convergence of proliferation-inducing pathways on p-S6 allows IL-21 to amplify the B cell response by increasing the basal p-S6 amount thereby facilitating B cell activation and proliferation in conjunction with BCR or CD40 signaling.

BCR signaling in naïve B cells is increased compared to GC B cells [53], especially through NF-kB activation [40, 54]. As a result, the strength of BCR signaling, determined by the affinity for antigen [10], can strongly influence B cell activation. The initiation of the cell cycle in naïve B cells *in vivo* is a multi-step process in which BCR signaling changes the metabolic state of the cell but is, by itself, insufficient to initiate the cell cycle [55] with the subsequent, timely receipt of T cell help inducing proliferation [55]. Antigen affinity may therefore regulate B cell response initiation via BCR signaling by determining the efficiency of antigen uptake and thus access to T cell help [12]. However, the magnitude and breadth of a B cell response is also related to the inflammatory stimuli delivered by vaccination or infection. The production of cytokines by CD4 T cells is one way by which information about the nature of an infection is conveyed to B cells, resulting, for example, in differential antibody isotype class-switch recombination [14, 56, 57]. The results presented here reveal an additional way by which T cells regulate B cell responses, namely IL-21-mediated modulation of naïve B cell proliferation. Our *in vitro* experiments, while not directly assessing BCR affinity-dependent signaling, imply that the *in vivo* consequences of IL-21 on early B cell activation are – at least in part – a result of the additive nature of BCR, CD40 and IL-21R on AKT and S6 phosphorylation. In support of this, p-S6 was found to be highly enriched in GC B cells expressing c-Myc [58], a population of cells about to enter cell division as a consequence of receiving T cell help [59].

IL-21 production by CD4 T cells is induced by IL-6 [60, 61] and calcium signaling via NFAT [62, 63] and thus is in response to inflammatory signaling with a STAT3-dependent autocrine loop stabilizing its production [64]. In light of its potent role in B cell activation reported here, this enables IL-21 to fine-tune B cell responses in relation to the immunological properties of the immunogen. Equally, excessive IL-21 production could result in the potentially detrimental lowering of BCR or CD40 signaling thresholds and/or exaggerated B cell expansion and thus predispose to autoimmunity. This could provide a mechanistic explanation for the association of polymorphisms in *Il21* and *Il21r* [65, 66] and increased IL-21 production [67] with systemic lupus erythematosus (SLE).

While IL-21 has long been reported to promote PC differentiation [17, 27, 47, 68, 69], this association is complicated by PC differentiation being tightly coupled to the extent of cell division [70]. Based on the close correlation of GC B cells and PC numbers irrespective of IL-21R expression revealed here, we now suggest that IL-21’s primary role in promoting antibody production and PC differentiation is by initiating and sustaining B cell proliferation rather than altering the balance in the bifurcation between GC B cell and PC fates. Although we only investigated the early stage of the GC response and did not address the role of IL-21 in the affinity-dependent selection of long-lived PC [34], it seems likely that IL-21 could also affect this process. We would expect this effect to be indirect, via its impact on proliferation changing both the size and duration of the GC response [27] and the extent of affinity maturation, which is also related to cell division [71]. Our findings that IL-21 increased the speed of passage through and frequency of entry into the cell cycle and promoted GC B cell accumulation over a large range of BCR affinities together with its previously reported roles in maintaining GC [27, 28] and the LZ/DZ ratio [29, 46] are all indicative of the key role of IL-21 in the initiation of a TD immune response being to promote the proliferation of pre-GC and GC B cells. We previously described the role of IL-21 in GC LZ B cell proliferation [29] while another recent study showed that the cyclic re-entry of LZ GC B cells could still occur when MHC-II or T cells had recently been deleted [72]. These results, with those presented here, suggest to us that the lingering presence of IL-21 may sustain GC B-cell proliferation following its initiation by cell contact mediated mitogenic signals.

In summary, by increasing both cell cycle initiation and speed, IL-21 modulates the breadth and magnitude of GC initiation and PC output. These results provide a novel mechanism by which IL-21 influences immune responses including those to vaccination and infection as well as a potential involvement in autoimmunity.

## Materials and methods

### Mice, cell transfer and immunization

SW_HEL_ mice (V_H_10_tar_IgH, Vκ10-κ Tg) [31] were crossed with *Rag1*^-/-^ (L. Corcoran, WEHI, Australia) and *Il21r*^-/-^ mice (W. Leonard, NIH, USA). OTII mice [32] (W. Heath, University of Melbourne, Australia) were crossed with *Il21 Gfp* knock-in mice [22] to obtain IL-21-GFP reporter mice (*Il21*^Gfp/+^) or IL-21 deficient mice (*Il21*^Gfp/Gfp^, referred to as *Il21*^*-/-*^). All mice were bred under specific pathogen free (SPF) conditions within the Monash Animal Research Platform and experimental mice were housed under SPF conditions within the Alfred Alliance Monash Intensive Care Unit. The ARA Animal Ethics Committee (Application E/1787/2018/M) approved all animal studies. Male and female mice were used throughout the study and experimental groups were matched based on age and gender. For adoptive cell transfer, spleens of OTII and SW_HEL_ mice were passed through a 70 µM mesh, red blood cells lysed, and the frequency of T and B cells determined by flow cytometry. Additionally, B cells were depleted from OTII splenocytes by magnetic sorting using CD45R (B220) MicroBeads according manufacturer’s instructions (Miltenyi Biotec cat. 130-049-501). For experiments involving cell division analysis, SW_HEL_ B cells were labelled with CTV (Thermo Fisher cat. C34557) according to the manufacturer’s instructions. Per recipient mouse, a mix of 1 × 10^5^ SW_HEL_ (50 % WT, 50 % *Il21r*^-/-^) and 5 × 10^4^ OTII T cells was then transferred *iv* and mice were immunized *ip* with 50 µg HEL^WT^OVA_pep_, HEL^2X^OVA_pep_ or HEL^3X^OVA_pep_ adsorbed on 45 or 90 µg alum adjuvant (Alhydrogel, InvivoGen cat. 21645-51-2).

### HEL-OVA_pep_ protein production

The nucleic acid sequence of HEL^WT^, HEL^2X^ (HEL with D101R and R73E mutations [73]) or HEL^3X^ (HEL with D101R, R73E and R21Q mutations [73]) fused to OVA_217-345_ and a deka-HIS tag was cloned into the pcDNA3.1 plasmid. Expi293 or HEK293E cells were transfected using polyethyleneimine (PEI) following culture for 5 days. TALON Superflow Metal Affinity Resin (Takarabio cat. 635506) was used to purify the recombinant HEL-OVA_pep_ protein. After dialysis against PBS, and concentration to 0.8-1.2 mg/mL (Amicon Ultra-15 Centrifugal Filter Units, Merck cat. UFC901024), the final protein was analyzed by polyacrylamide gel electrophoresis and Coomassie blue staining, aliquoted and frozen at - 80°C.

### Flow cytometry

Spleens were isolated and passed through a 70µM mesh to generate a single cell suspension. Following red-blood cell lysis, up to 5 × 10^7^ cells were stained with monoclonal antibodies (Table 1) to cell surface proteins diluted in PBS containing 1 % BSA (Bovogen) and 0.1 % NaN_3_ (Sigma) (staining buffer) and in the presence of FcγR blocking antibody (clone 2.4G2, WEHI Antibody Facility) and 1 % rat serum on ice for 30 minutes. In experiments where antigen-binding was analyzed, cells were first incubated with 400 ng/mL biotinylated HEL^WT^OVA_pep_, HEL^2X^OVA_pep_ or HEL^3X^OVA_pep_ in staining buffer on ice for 30 minutes, then washed once and then incubated with monoclonal antibodies and fluorochrome-conjugated streptavidin. Dead cells were excluded using Fixable Viability Dye eFluor™ 780 (eBioscience, cat. 65-0865-14) or FluoroGold (Santa Cruz Biotechnology, CAS 223769-64-0). Cells were analyzed using BD LSR Fortessa X-20, BD LSR-II or Cytek Aurora flow cytometers. The data were analyzed with FlowJo (BD) and SpectroFlo (Cytek) software and statistical analysis was performed using Prism 8 (GraphPad).

**Table 1:** Monoclonal antibodies, lectins and streptavidins used for flow cytometry.

| Antibody or lectin | Source | Identifier |
| --- | --- | --- |
| Active Caspase 3-BV650 (clone C92-605) | BD Biosciences | 564096 |
| B220-BV421 (clone RA3-B62) | BD Biosciences | 562922 |
| B220-PerCP-Cy5.5 (clone RA3-B62) | BD Biosciences | 551960 |
| BrdU-AF647 (clone 3D4) | BD Biosciences | 560209 |
| CD16/32 (clone 2.4G2) | WEHI Antibody Facility | N/A |
| CD4-BV510 (clone RM4-5) | BioLegend | 100559 |
| CD4-A680 (clone GK1.5) | WEHI Antibody Facility | N/A |
| CD4-PerCP-Cy5.5 (clone RM4-5) | BD Biosciences | 550954 |
| CD19-BUV737 (clone 1D3) | BD Biosciences | 612781 |
| CD45.1-APC-eFluor780 (clone A20) | Invitrogen | 47053-82 |
| CD45.1- PerCP-Cy5.5 (clone A20) | eBioscience | 45-0453-80 |
| CD45.2-BV786 (clone 104) | BD Biosciences | 563686 |
| CD98-BV711 | BD Biosciences | 745466 |
| CD98-PE (clone RL388) | BioLegend | 128208 |
| CD138-BV650 (clone281-2) | BD Biosciences | 564068 |
| CXCR5-Biotin (clone 2G8) | BD Biosciences | 551960 |
| FAS-BUV395 (clone Jo2) | BD Biosciences | 740254 |
| GL7-PE (clone GL7) | BD Biosciences | 561530 |
| IgD-BV711 (clone 11-26c.2a) | BD Biosciences | 564275 |
| IgD-BV421 (clone 11-26c.2a) | BD Biosciences | 744291 |
| p-AKT (S473)-PE (clone M89-61) | BD Biosciences | 561671 |
| PD-1-PE (clone J43) | BD Biosciences | 551892 |
| PNA-FITC | Vector Laboratories | FL-1071 |
| p-S6 (Ser235/236)-PE-Cy7 (clone D57.2.2E) | Cell Signaling | 34411S |
| Streptavidin-A647 | Invitrogen | 84E2-1 |
| Streptavidin-PE-Cy7 | eBioscience | 25-4317-82 |
| Streptavidin-BV650 | BD Biosciences | 563855 |
| Streptavidin-BV786 | BD Biosciences | 563858 |
| TCR-Vα2-APC (clone B20.1) | WEHI Antibody Facility | N/A |
| TCR-Vβ5.1, 5.2-PE-Cy7 (clone MR9-4) | BioLegend | 139508 |

### BrdU incorporation and 7AAD staining

For *in vivo* BrdU incorporation, mice were injected *ip* with 200 µL of 10 mg/mL 5-bromo-2’-deoxyuridine (BrdU, BD cat. 559619). Mice were culled by cervical dislocation at the time point indicated in the individual experimental setup, spleens were harvested, and 6 × 10^7^ cells stained with monoclonal antibodies against cell-surface molecules as described above. After washing with FACS buffer, cells were suspended in 200 µL BD Cytofix/Cytoperm (BD cat. 554722) and incubated for 15 min on ice followed by washing with 2 mL BD Perm/Wash. If staining for active caspase-3, the cells were then incubated for 30 min with anti-active caspase 3 in BD Perm/Wash (BD cat. 554723). Next, cells were suspended in 150 µL BD Permeabilization Buffer Plus (BD cat. 561651) on ice for 10 min followed by washing with 2 mL BD Perm/Wash. Cells were fixed again with 150 µL BD Cytofix/Cytoperm on ice for 5 min followed by washing with 2 mL BD Perm/Wash. To increase BrdU accessibility for monoclonal antibodies, DNAse digest was performed. Per sample a mix of 60 µL DNAse1 stock (1 mg/mL in ddH_2_O, Sigma, Cat. D4513) and 140 µL PBS was freshly prepared and cells were incubated at 37°C for 1 h. After washing with 1 mL BD Perm/Wash, cells were incubated at room temperature with fluorochrome labelled anti-BrdU diluted in BD Perm/Wash. After a final wash with 2 mL BD Perm/Wash, cells were either resuspended in 1 mL FACS buffer or incubated with 20 µL 7-AAD for 5 minutes followed by the addition of 1 mL FACS buffer and acquired as described above.

### In vitro cell culture, stimulation and phosphoflow staining

For phosphoflow analysis, splenocytes were isolated from male or female C57Bl/6 and *Il21r*^-/-^ mice and 3 × 10^6^ cells used per stimulation condition. Cells were incubated for 3 hours at 37°C in RPMI media supplemented with or without recombinant mouse IL-21 (20 ng/mL, Peprotech cat. 210-21-100). Cells were stimulated with either anti-Igκ and anti-Igλ or anti-CD40. For anti-Igκ and anti-Igλ stimulation, biotinylated rat anti-Igκ (100 ng/mL; clone 187.1, WEHI Antibody Facility) and anti-Igλ (100 ng/mL; clone JC5, WEHI Antibody Facility) were added during the final 15 min of the 3h incubation, followed by addition of avidin (10 µg/mL) for 1 min. For anti-CD40 stimulation, cells were incubated with anti-CD40 (20 ng/mL; clone 1C10, WEHI Antibody Facility) throughout the 3h culture period. Stimulation was stopped by fixation and permeabilization with the BD phosphoflow staining reagents (BD cat. 558049 and 558050) as per manufacturer’s instructions. Cells were stained for flow cytometry with anti-p-AKT, anti-p-S6, anti-B220, anti-IgD and PNA (see Table 1 for further details). Flow cytometry was performed on a LSRFortessa X20 flow cytometer (BD) or a Cytek Aurora (experiments shown in Fig. 5 and 6). Flow cytometry data were analyzed with FlowJo 10 software (BD). For *in vitro* cell culture of SW_HEL_ B cells, 1 × 10^6^ splenocytes from a RAG^-/-^ SW_HEL_ mouse were incubated per well of a 96-well U-bottom plate in RPMI media supplemented with 5 % fetal calf serum, 1 µg/mL HEL^WT^OVA_pep_, IL-4 (10 ng/mL) and additional stimuli as indicated.

### Statistical analysis

All statistical analyses were performed using Prism 8 (GraphPad). Mice failing to respond to immunization, as evidenced by failure to expand adoptively transferred B cells, were excluded from analysis. No blinding was done. Absolute cell numbers were calculated based on total spleen count and proportional representation in flow cytometry with electronic gates containing < 5 events excluded from analysis. In experiments where the expansion of WT and *Il21r*^-/-^ B cells was studied within the same animal, cell counts were corrected for any deviation of a 1:1 ratio at the time of transfer.

## Acknowledgements

We thank Lynn M. Corcoran, William R. Heath, Warren J. Leonard and Stephen L. Nutt for providing mouse strains. We thank the Alfred Alliance Monash Intensive Care Unit, and Monash Animal Research Platform for animal husbandry and Stephanie Jansen for assistance with intravenous injections. The authors acknowledge the contributions of AMREPflow, ARAFlowCore, Noelene Quinsey from the Monash Protein Production Unit; Tim Adams, Tam Pham, Tram Phan and George Lovercz from the Commonwealth Scientific and Industrial Research Organisation (CSIRO). A.R.D was supported by a Monash University Research Training Program (RTP) Stipend and Z.D. by a Swedish International Postdoctoral Fellowship (2016-06659). D.M.T. was funded by National Health and Medical Research Council (NHMRC) Australia Investigator Award (APP1175411), I.Q. by an Early Postdoc Mobility fellowship (P2ZHP3_164964) and an Advanced Postdoc Mobility fellowship (P300PA_177893) provided by the Swiss National Science Foundation and a Peter Doherty Early Career fellowship (APP1145136) provided by NHMRC Australia. This work was supported by NHMRC Project Grant APP1146617 awarded to D.M.T, D.Z and I.Q, Monash Platform Access grants PAG17-0207 awarded to D.Z, I.Q and D.M.T and PAG18-0409 awarded to I.Q. and D.Z, NHMRC Ideas Grants APP1185294 awarded to M.J.R and I.Q. and APP2002393 awarded to I.Q.

## Author contributions

Conceptualization: DMT, IQ; Methodology: ARD, CIM, RB, DZ, IQ; Investigation: ARD, CIM, MJR, ZD, CP, KOD, IQ; Writing- Original Draft: IQ; Writing- Review and Editing: ARD, CIM, MJR, ZD, DZ, RB, DMT, IQ.

## Extended view figure legends

**Fig. EV1: SW**_**HEL**_ **B cells characterization and flow cytometry gating strategy for Fig. 1-3**.

**(A, B)** Lymphocytes from SW_HEL_ mouse spleen. **(C)** HEL-OVA_pep_ variant binding to SW_HEL_ B cells. **(D)** Exemplary electronic gating strategy for Fig. 1B-E to identify WT (GFP^-^) and *Il21r*^-/-^ (GFP^+^) SW_HEL_ B cells. **(E)** Gating for Fig. 2B, 2D and 3C. SW_HEL_ B cells were identified by CTV and HEL^2X^OVA_pep_ antigen-binding. eGFP was used to distinguish WT and *Il21r* ^-/-^ cells. Blue gates were used for analysis in Fig. 1I and red gates for Fig. 3C.

**Fig. EV2: Flow cytometry gating strategy for Fig. 4**.

Exemplary sequential electronic gating strategy as applied in Fig. 4 to identify naïve B cells following *in vitro* culture and phosphoflow staining.

**Fig. EV3: Flow cytometry gating strategy for Fig. 5**.

**(A)** Exemplary electronic gating strategy for Fig. 5B-C. OTII T cells were identified based on their co-expression of TCR Vα2 and β5 and Tfh by co-expression of PD-1 and CXCR5. **(B)** Gating strategy for Fig. 5D-G. Gates 1-4 were activated in SpectroFlo software after which CD45.2 positive cells were exported for analysis using FlowJo. **(C)** For proliferation analysis by CTV dilution, WT or *Il21r*^-/-^ SW_HEL_ B cells from individual mice were concatenated into one file per experiment. CTV division peaks were then identified using FlowJo’s Proliferation tool by first identifying peaks 0-7 and then 8-11+. Gray bars indicate gate used to set the initial peak (0 or 4). For statistical analysis in Figure 5E, resultant electronic gates for each CTV division were then applied to individual mice.

**Fig. EV4: Flow cytometry gating strategy and additional data for Fig. 6**.

**(A)** Gating strategy for Fig. 6. Initial gating in SpectroFlo software as shown in Figure EV3, after which CD45.2 positive cells were exported for subsequent analysis in FlowJo. Red gates indicate gates to identify plasma cells. **(B)** Exemplary expression of Fas and GL7 on total SW_HEL_ B cells. **(C)** Exemplary CD86 and CXCR4 expression on WT or *Il21r*^-/-^ SW_HEL_ GC B cells showing gates to identify LZ cells and **(D)** statistical analysis by Multiple paired t tests with p-values corrected for multiple comparisons using Holm-Šídák method. **** p ≤ 0.0001. Data were pooled from two independent experiments.

