## Supplementary figures and images for "IL-21 has a critical role in establishing germinal centers by amplifying early B cell proliferation"

### Supplemental Figures 1-4

#### Extenden View Figure 1

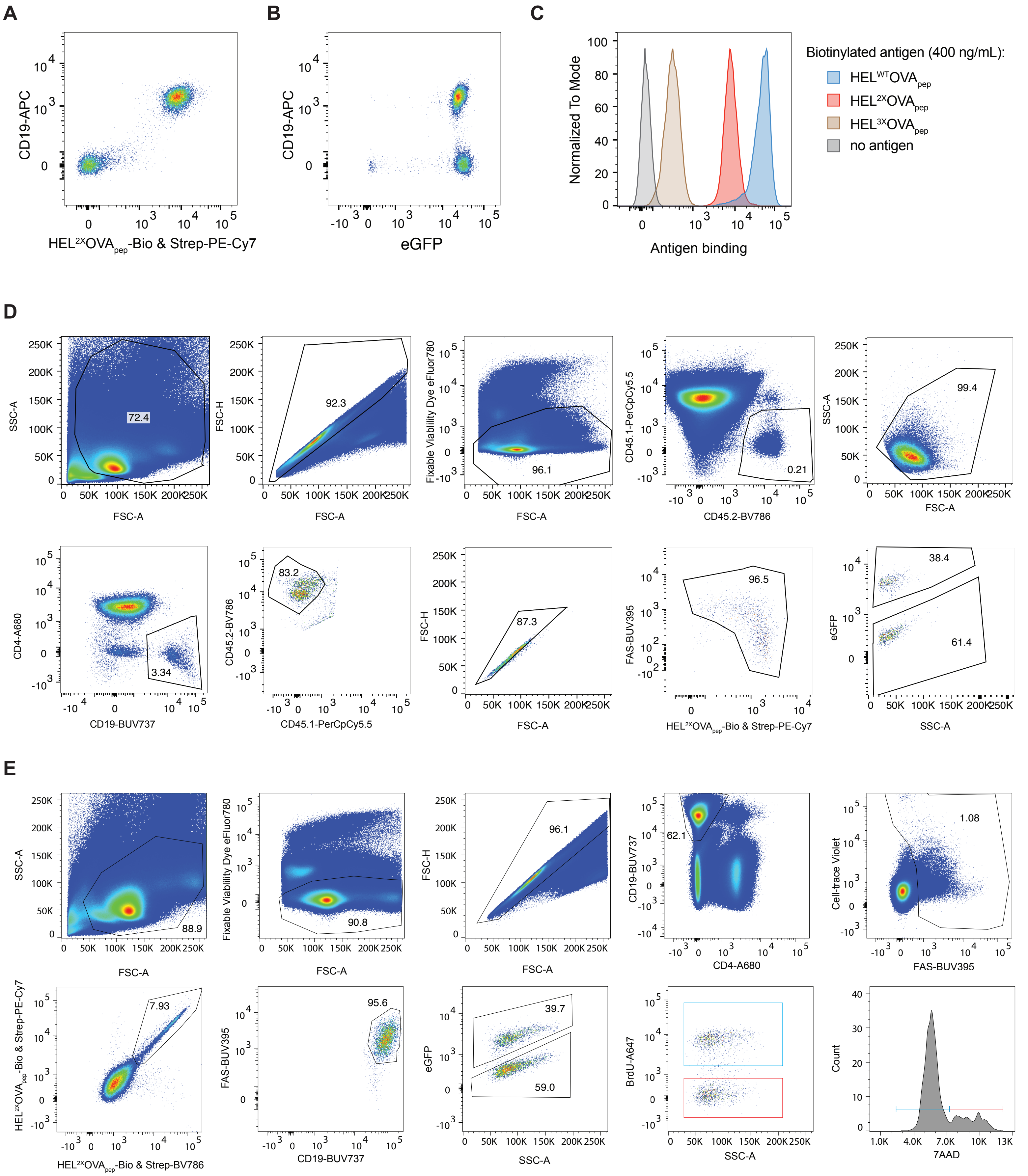

#### Extenden View Figure 2

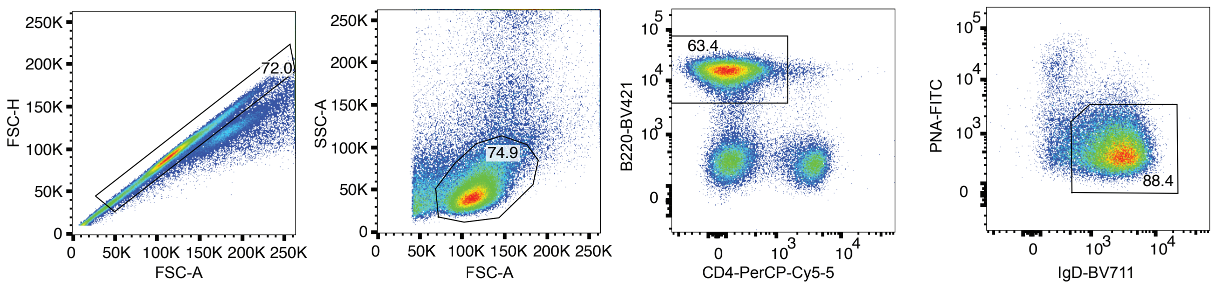

#### Extenden View Figure 3

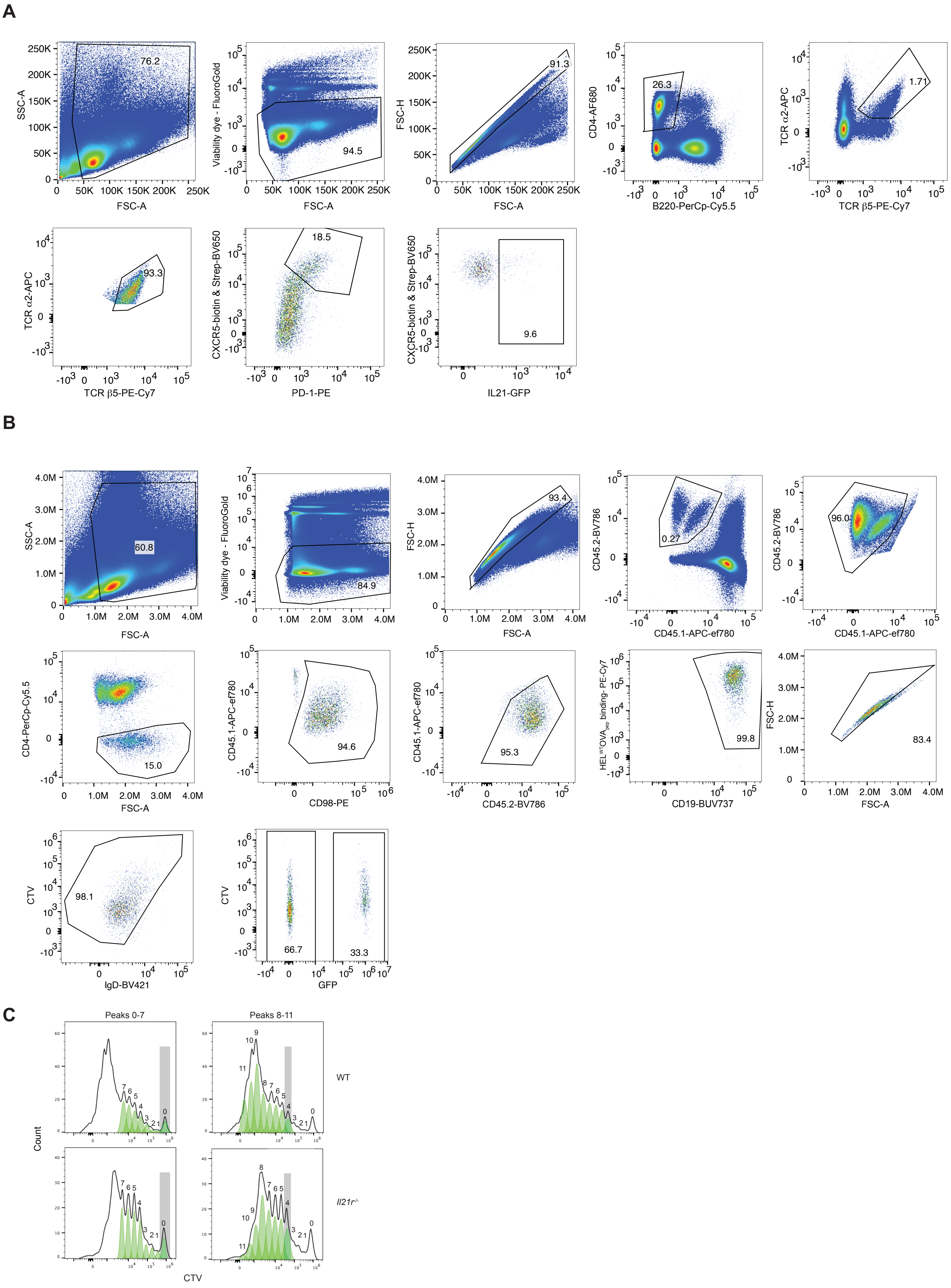

Extenden View Figure 4

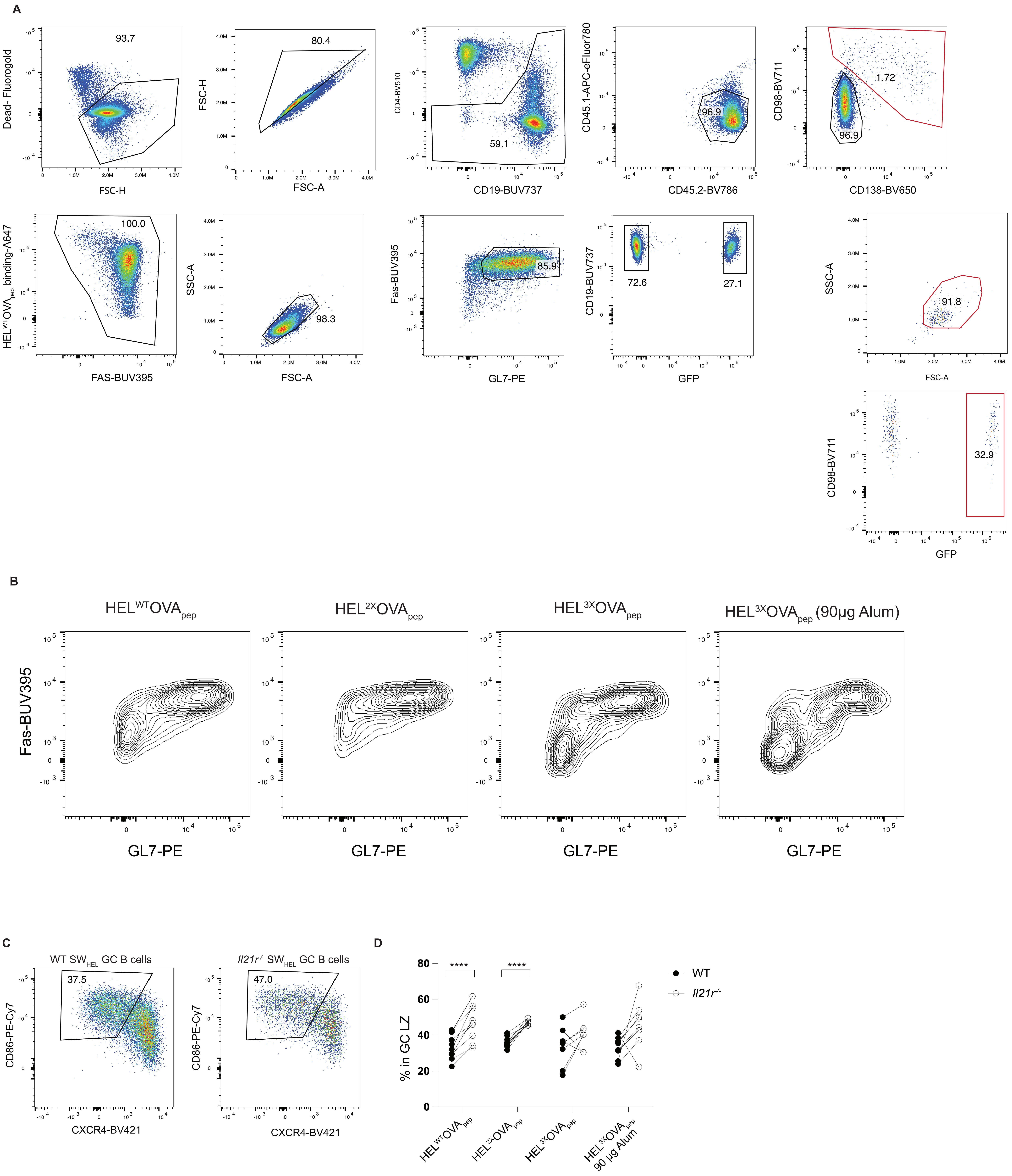
